# AIPAMPDS: an AI platform for antimicrobial peptide design and screening

**DOI:** 10.1101/2025.03.24.644774

**Authors:** Yanru Li, Liming Gao, Wenqian Zhao, Ziyu Wang, Xianghan Xu, Jinwen Ji, Yuan Zhang, Yufeng Liu, Liangyun Zhang, Yong Yang, Jinhu Huang, Cong Pian

**Author notes:** To whom correspondence should be addressed. Correspondence may also be addressed to Jinhu Huang. Correspondence may also be addressed to Yong Yang. The first three authors should be regarded as Joint First Authors.

## Abstract

Antimicrobial resistance (AMR) presents a pressing global health crisis, particularly in the post-pandemic era, underscoring the urgent need for innovative therapeutics. Antimicrobial peptides (AMPs) hold promises for combating multidrug-resistant infections but are hindered by challenges in clinical translation, including limited hemocompatibility and inefficient design processes. To overcome these obstacles, we developed AIPAMPDS, an AI-driven computational platform that integrates a deep generative model with multi-stage screening to design highly active, non-haemolytic AMPs. The platform features two key modules: (1) a Generative Pre-trained Transformer (GPT)-based framework for target-specific AMP generation, enabling rapid exploration of novel peptide sequences, and (2) a multi-neural network screening pipeline for dual evaluation of antimicrobial activity and toxicity. By systematically evaluating both AI-generated and natural candidate AMPs, we identified ten lead peptides for experimental validation, of which 90% exhibited negligible haemolytic activity while maintaining robust antimicrobial potency. AIPAMPDS bridges computational design and therapeutic application, significantly advancing AMP discovery and providing a powerful tool for next-generation antimicrobial development. The platform is freely accessible via a web server at https://amps.pianlab.cn/AIPAMPDS/.

**GRAPHICAL ABSTRACT:** 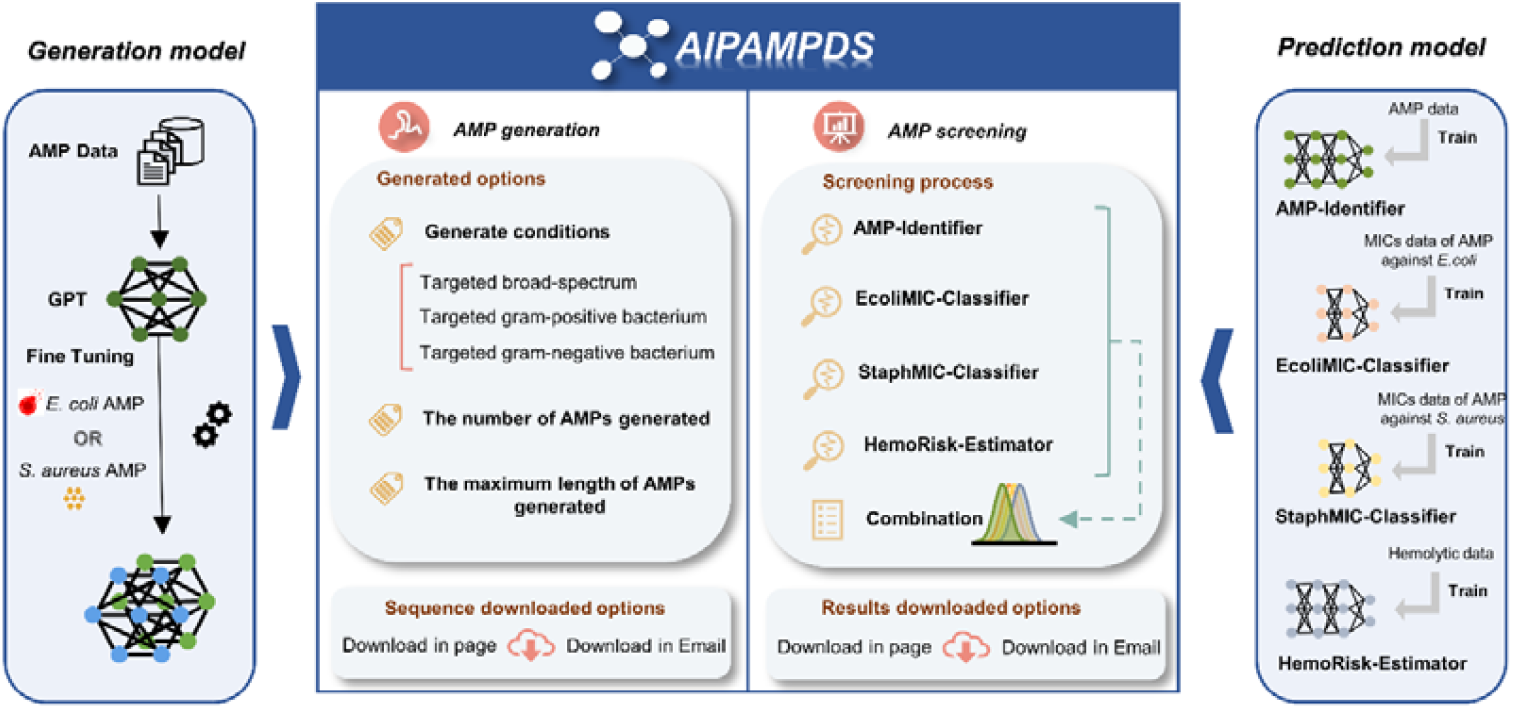

## INTRODUCTION

The escalating overuse and misuse of antibiotics have precipitated a global surge in multidrug-resistant pathogens, posing an imminent health crisis that undermines the efficacy of conventional therapies and necessitates innovative antimicrobial strategies(1). Antimicrobial peptides (AMPs) have emerged as promising candidates for anti-infective drugs due to their ability to combat infections caused by bacteria, fungi, viruses, and other microorganisms(2-4). Unlike conventional antibiotics, AMPs exhibit distinct pharmacodynamic properties that reduce the likelihood of resistance development(5). Typically comprising 10–50 amino acids (AAs), AMPs exert their antimicrobial effects by targeting bacterial cell membranes, forming transmembrane ion channels, and disrupting membrane integrity, which leads to the leakage of cellular contents and ultimately bacterial cell death(6).

Artificial intelligence (AI) and machine learning (ML) methodologies, particularly statistical learning frameworks and optimization-based approaches, have shown considerable potential in designing both small molecules and macromolecules, including AMPs. The integration of AI into this field encompasses multifaceted approaches, incorporating generative models and classification techniques(7). Deep generative models have emerged as promising computational tools for therapeutic peptide design(8), capable of capturing the underlying data distribution of a given dataset to generate novel instances that faithfully reflect the properties of the original data(9,10). Current implementations predominantly utilize advanced deep learning architectures such as Generative Adversarial Networks (GANs)(11,12) and Variational Autoencoders (VAEs)(13-15), along with conditional variants like seqGAN(16) and cVAEs(7,17,18). Concurrently, a diverse array of computational techniques has been developed for AMP discovery(19). Traditional ML-based approaches have identified AMPs by employing Support Vector Machine (SVM)(20-22), Random Forest (RF)(21-25) and ensemble learning methods(26). Deep learning (DL)-based methods, such as Convolutional Neural Networks (CNNs)(27), Long Short-Term Memory (LSTM) networks, Graph Attention Networks (GAT)(28), and their combinations(29,30), enable automated identification of functional peptides. Notably, pretrained protein language model-based methods(16,31-35) used the ESM-2 protein language model(36) and ProtTran(37), which are widely used for deep learning tasks involving protein sequence data.

Despite these advances, significant limitations persist. AMP generation models like GANs and VAEs struggle to capture long-range sequence correlations, often yielding peptides with reduced diversity and authenticity(38). Specifically, GANs are susceptible to training instability and mode collapse, while VAEs face challenges in generating diverse sequences due to spatial distribution constraints(39,40). Furthermore, existing screening frameworks rarely optimize both antimicrobial activity and hemolytic toxicity concurrently during AMP design. Critically, comprehensive computational platforms that integrate AMP generation and screening are scarce.

To address these challenges, we developed AIPAMPDS (AI Platform for Antimicrobial Peptide Design and Screening), a unified system featuring two synergistic modules: (1) a Transformer-based generation module for de novo AMP synthesis, and (2) a multimodal screening module that integrates BERT-based classification of antimicrobial activity, CNN-driven prediction of minimum inhibitory concentration (MIC), and RNN-based hemolysis prediction. To enhance accessibility, AIPAMPDS is available as both a web-based platform (https://amps.pianlab.cn/AIPAMPDS/) and a local-server implementation, supporting user-configurable deployment across diverse computing environments.

## MATERIAL AND METHODS

### Data selection and processing

Six datasets were curated for this study: AMP, non-AMP, MIC, fine-tuning, pre-trained AMP, and hemolytic datasets **(Fig. 1A)**. Details of their preparation are as follows:

**Figure 1.**
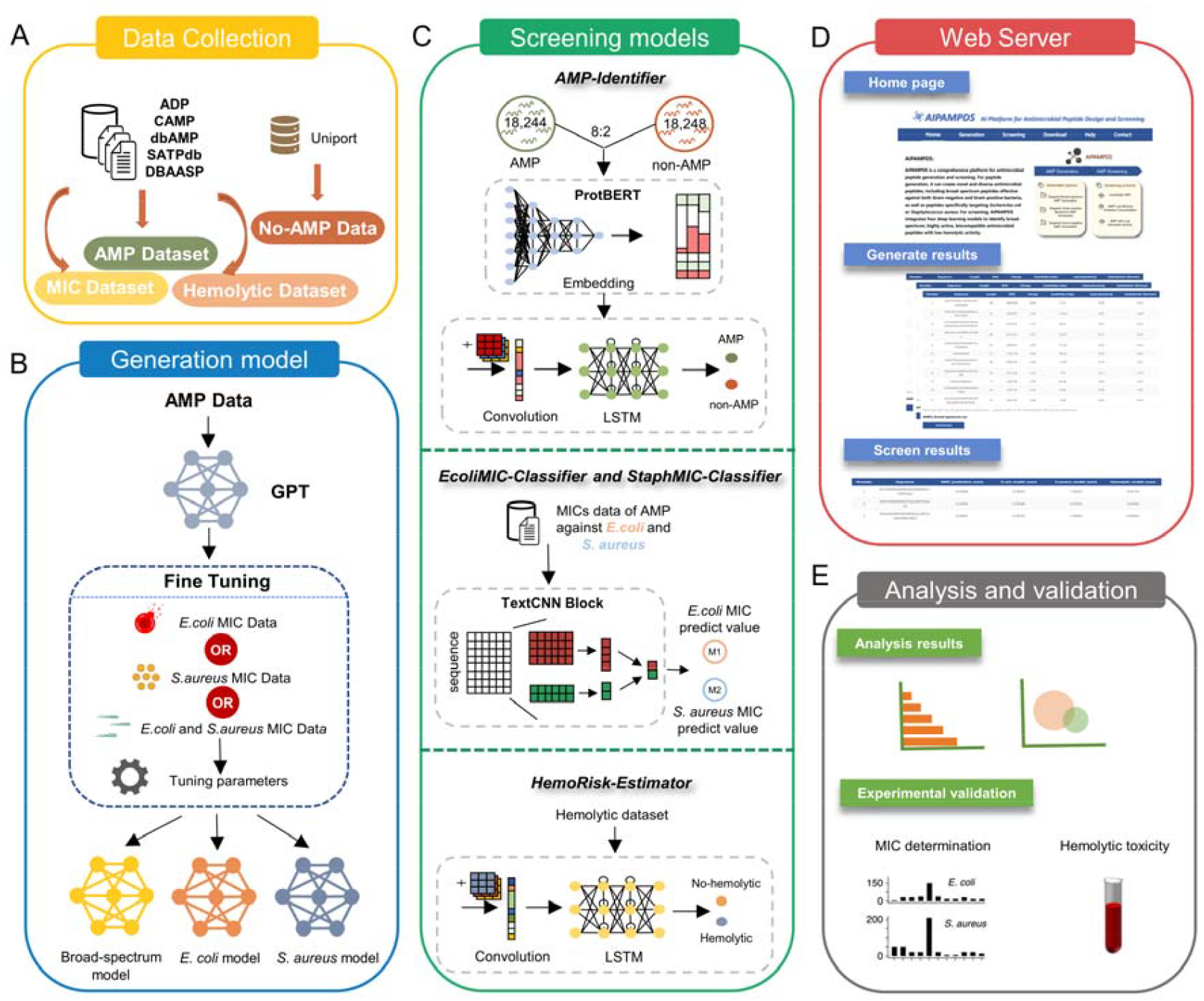
The procedure of the development of AIPAMPDS, including data collection, generation model construction, screening model construction, web server, analysis and experimental validation. Collection and pre-processing of AMPs from the public databases. **(B)** AMP generation model: training of generative models using GPT. The three generative models obtained are broad-spectrum AMPs generation model, against *E. coli* AMPs generation model and *S. aureus* generation model. (C) Screening models: four machine learning approaches, including transformer, DNN, CNN, LSTM and attention were used for the construction of predictive models (AMP-Identifier, EcoliMIC-Classifier、StaphMIC-Classifier and HemoRisk-Estimator). (**D**) Screenshot of the web page. (**E**) Analysing results as well as experimental verification of MIC values and hemolysis.

### AMP dataset

AMPs were retrieved from the five major public AMP databases, including dbAMP(41), APD(42), CAMP(43), SATPdb(44) and DBAASP(45)—encompassing a broad range of sources (downloaded as of April 2023). After removing duplicates, 31,552 sequences (3–300 AAs) were compiled. Sequences containing ambiguous “X” residues were excluded, yielding 28,508 sequences are obtained as the data of the AMP classification model.

### Non-AMP dataset

Non-AMP sequences were sourced from UniProt(46) (http://www.uniprot.org) by filtering for cytoplasmic peptides and excluding entries with keywords “antibiotic,” “antimicrobial,” “antifungal,” “antiviral,” “effector,” or “excreted” (downloaded April 2023). Duplicates and sequences exceeding 300 AA were removed, as were those matching the AMP dataset. The final set, matched to the AMP dataset’s length distribution, comprised 28,513 sequences.

### Fine-tuning dataset

MIC data for fine-tuning the generative model of *Escherichia coli*(*E. coli*) and *Staphylococcus aureus*(*S. aureus*) was obtained from dbAMP(41) (downloaded April 2023). Sequences with MIC values listed in the “targets” column were recorded, with units converted to µg/ml. After removing duplicates and sequences containing “X”, the dataset yielded 3,106 sequences for *E. coli*, 5,266 for *S. aureus*, and 6,239 for combined *E. coli* and *S. aureus* fine-tuning.

### Pre-trained AMP dataset

This dataset was derived from the AMP dataset by excluding sequences that were duplicated in the fine-tuning *E. coli* and *S. aureus* MIC datasets. Sequences shorter than 5 AAs or longer than 190 AAs were also removed, resulting in a total of 22,122 AMPs.

### MIC dataset

AMP sequences annotated with MIC values were collected from DBAASP(45), dbAMP(41), CAMP(43), and DRAMP(47) (downloaded April 2023) and merged into an integrated dataset with duplicates eliminated. Sequences specific to *E. coli* and *S. aureus* with MIC values were extracted, excluding those with non-natural AAs. Only sequences of 5–50 AAs were retained, and MIC values were standardized to µg/ml. Sequences with MIC ≤16 µg/ml were classified as positive (highly active), and those >16 µg/ml as negative. The final dataset included 4,258 positive and 4,670 negative sequences for *E. coli*, and 3,953 positive and 4,496 negative sequences for *S. aureus*.

### Hemolytic dataset

Hemolysis data were sourced from DBAASP(45), which provides peptides annotated with hemolytic activity. Peptides causing <20% hemolysis at concentrations ≥50 mM were classified as non-hemolytic, while those causing >20% hemolysis at any concentration were deemed hemolytic, yielding 2,438 hemolytic and 1,539 non-hemolytic sequences.

### Dataset Splitting

The AMP and non-AMP datasets were divided into training-validation and test datasets at an 8:2 ratio. The training-validation set was further split into training and validation subsets, also at an 8:2 ratio. The training dataset (18,244 AMPs, 18,248 non-AMPs) was used for model training, the validation dataset (4,562 AMPs, 4,562 non-AMPs) for hyperparameter tuning, and the test dataset (5,702 AMPs, 5,703 non-AMPs) for performance evaluation against existing models.

### Architecture design

The AIPAMPDS platform was constructed to facilitate AMP generation and screening, leveraging five key deep learning models: the AMP Generation Model, AMP-Identifier, EcoliMIC-Classifier, StaphMIC-Classifier, and HemoRisk-Estimator.

### AMP generation model

The AMP generation model was a mini version of the Generative Pre-Training Transformer (GPT) model(48), which focused on predicting the next AA through masked self-attention to generate drug-like peptides (**Fig. 1B**). It employs a transformer decoder comprising eight stacked blocks, each featuring a masked self-attention layer followed by a fully connected feed-forward neural network. The self-attention layer outputs a 256-dimensional vector, which serves as input to the feed-forward network. Within each block, a hidden layer expands this to a 1,024-dimensional vector, applies GELU activation layer, and the final layer of the fully connected neural networks reduces it back to 256 dimensions for the subsequent block. Training proceeded in two phases: pre-training on the pre-trained AMP dataset for 30 epochs with a learning rate of 6 × 10□□, followed by fine-tuning on the fine-tuning dataset for 30 epochs at a learning rate of 1 × 10□□. AdamW(49) was used in the optimizer, β□ = 0.9, β□ = 0.95. For sequence generation, the model initiates with a starting token and autoregressively predicts subsequent AAs, ultimately producing 100,000 peptide sequences.

### AMP-Identifier

We developed the AMP-Identifier to classify AMPs versus non-AMPs, building upon a natural language processing (NLP)-inspired deep learning framework **(Fig. 1C)**. This model adapts ProtBERT(37), a transformer-based architecture pre-trained on approximately 217 million protein sequences from the Big Fantastic Database (BFD)(50,51), featuring 12 attention heads and 12 hidden layers. ProtBERT(37) processes peptide sequences by treating AAs as words and peptides as sentences, employing sinusoidal positional encoding and multi-head self-attention to capture token dependencies. While ProtBERT(37) provides robust sequence embeddings through its self-attention mechanism—computing weighted sums of values (V) based on query (Q) and key (K) dot products normalized by softmax—standard fine-tuning typically applies global average pooling, potentially discarding fine-grained sequence details critical for AMP classification.

To address this limitation and enhance performance, we retained all outputs for the subsequent layer and extended ProtBERT(37) with a custom architecture tailored for AMP identification. The AMP-Identifier architecture incorporates the ProtBERT(37) model for embedding sequences, a convolutional layer, a 1D max pooling layer (pool_size: 5, strides: 5), an LSTM layer (units: 100, unroll: True, stateful: False), and a dense layer. For binary classification (AMP: 1, non-AMP: 0), a fully connected layer with sigmoid activation generates probabilities, with a stringent threshold of 0.998 to define AMPs, ensuring high specificity. The model was trained using the AdamW optimizer(49) with binary cross-entropy loss, an initial learning rate of 1 × 10□□, and a batch size of 32 for 12 epochs. The ReduceLROnPlateau scheduler reduced the learning rate by a factor of 0.1 if validation accuracy did not improve over four epochs. Various tuning parameters, such as learning rate and batch size, were considered, and hyperparameters were optimized based on accuracy (ACC).

### EcoliMIC-Classifier and StaphMIC-Classifier

To identify AMPs with high antimicrobial activity, we developed the EcoliMIC-Classifier and StaphMIC-Classifier, leveraging deep learning models to predict MIC **(Fig. 1C)**. Each model comprises three core components: an embedding module, a Text Convolutional Neural Network (TextCNN) module(52), and a classification module. During preprocessing, peptide sequences were converted into fixed-length encoding matrices (50 AAs) via numerical encoding and padding. The embedding module, with parameters set to input_dim: 21, output_dim: 192, and input_length: 50. The TextCNN(52) module, a multi-scale CNN, applies convolution operations with kernel sizes of 2 and 4 to the embedding matrix, capturing local AA correlations at varying scales. These kernel sizes were chosen to accommodate the minimum sequence length of 5 AAs in the MIC dataset, minimizing information loss during convolution. Post-convolution, feature maps undergo 1D max-pooling (pool size: 5) for dimensionality reduction, yielding a pooled representation that integrates global sequence features. This is followed by three fully connected layers, culminating in a sigmoid activation layer that outputs classification probabilities.

To achieve a specificity of 98%, we optimized the classification threshold by analyzing the false positive rate (FPR) on the test set, setting it to 0.998 to maintain an FPR below 0.02. This stringent threshold enhances the precision of identifying highly active AMPs by minimizing false positives. Both models were trained using the Adam optimizer with an initial learning rate of 1.8 × 10□^3^ and binary cross-entropy loss, a standard choice for binary classification. To improve convergence and adapt to training dynamics, a Cosine Annealing Scheduler dynamically adjusted the learning rate, gradually reducing it in later epochs to prevent overshooting and stabilize optimization.

### HemoRisk-Estimator

The HemoRisk-Estimator was designed to predict hemolysis risk by modeling position-invariant sequence patterns in AMPs, using a deep neural network (DNN) implemented via the Keras framework (http://keras.io) with TensorFlow(53) as the backend **(Fig. 1C)**. The architecture integrates convolutional layers, max-pooling layers, and an LSTM layer to capture both local and sequential dependencies in AMP sequences. Specifically, 1D convolutional layers generate filters to extract generalized sequence motifs, while max-pooling reduces spatial dimensions, and the LSTM layer models complex temporal patterns across diverse AMPs.

Drawing inspiration from Veltri *et al*.(29), we adopted a convolutional-LSTM-attention architecture for the construction of hemolysis prediction. Details of this architecture were as follows: the embedding layer (input_dim: 21, output_dim: 128 and input_length: 200); the 1D Conv layer (nb_filter: 64, filter_length: 64, strides: 1, activation: relu); 1D max pooling layer (pool_size: 5, strides: 5); LSTM layer (units: 100, unroll: True, stateful: False); Attention layer and dense layer (units: 1, activation: sigmoid). This design effectively correlates sequence patterns with hemolytic activity, providing a robust tool for AMP safety assessment.

### Architecture design

We used general quantitative indicators: ACC, precision (PRE), sensitivity (SEN), specificity (SPE), matthews correlation coefficient (MCC), F1-Score, area under the curve (AUC) of receiver operating characteristic (ROC) curve and AUC of the precision–recall (PR) curve to evaluate our model’s performance.

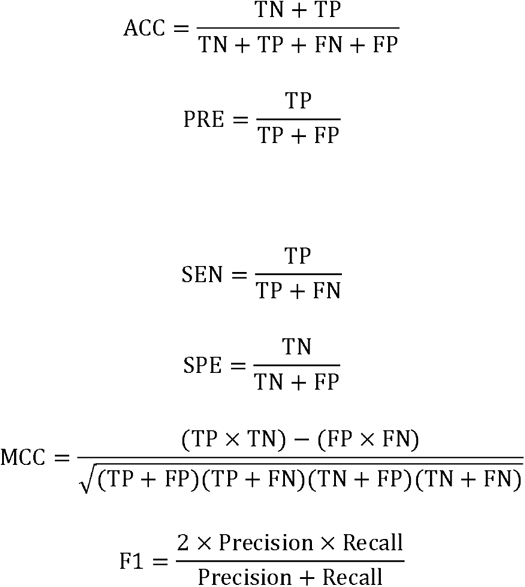

where TP, TN, FP and FN denote the number of true positives, true negatives, false positives and false negatives, respectively.

### Bacterial strains and growth conditions

All Gram-negative and Gram-positive bacterial strains were cultured in Luria-Bertani (LB) or trypticase soy broth (TSB) agar plates, respectively, and incubated overnight at 37 °C unless specifically indicated. A single colony of the tested bacterial strains was inoculated into medium broth with shaking

### Minimal inhibitory concentrations

Antimicrobial susceptibility testing was performed by broth microdilution method for AMPs against Gram-positive and Gram-negative pathogens above mentioned, following the methods of document CLSI M100-ED34(54). The Minimal inhibitory concentrations (MICs) were defined as the lowest concentrations that inhibit visible growth of the tested bacteria. *E. coli* ATCC 25922 and *S. aureus* ATCC 29213 were served as quality controls.

### Hemolysis assays

Red blood cells (RBCs) from Kunming mice were used to evaluate the hemolytic toxicity of the selected AMPs. Fresh red blood cells were collected from mice and washed until the supernatant was completely cleared. Added 19-fold volume of PBS to prepare 2% RBC suspension. Then 50 μL of RBC suspension was added to 50 μL of peptides at various concentration (1×MIC-16×MIC), followed by incubation at 37□ or 1 h. Afterwards, the mixture was centrifuged at 1500 rpm for 10 min. Then transferred 30 µL supernatant to 96-well plate containing 70 µL PBS. The absorbance of the sample was measured at 450 nm by a microplate reader. PBS and 2% Triton X-100 was used as negative and positive control, respectively. The experiment was performed in triplicate and the hemolysis activity was calculated by the formula:

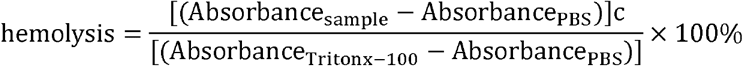

### The workflow of AIPAMPDS

AIPAMPDS serves as a comprehensive platform for the generation and screening of antimicrobial peptides (AMPs), with its workflow illustrated in **Fig. 1**. The generation pipeline produces novel and diverse AMP sequences, encompassing three categories: peptides targeting *E. coli*, peptides targeting *S. aureus*, and broad-spectrum peptides effective against both Gram-negative and Gram-positive bacteria. The screening pipeline processes input sequences sequentially through four integrated models—AMP-Identifier, EcoliMIC-Classifier, StaphMIC-Classifier, and HemoRisk-Estimator—to achieve a multi-step evaluation. This workflow first identifies AMPs, then filters for those with low minimum inhibitory concentrations (MICs), and finally selects peptides with minimal hemolytic activity. The detailed framework of each model is depicted in **Fig. 1**.

### Web server implementation

AIPAMPDS is deployed as a freely accessible web server at https://amps.pianlab.cn/AIPAMPDS/, hosted on a CentOS 7.6 system powered by an NVIDIA Tesla T4 GPU cluster (16 GB). The frontend is built with Vue 3 (https://vuejs.org/) and Element Plus (https://element-plus.gitee.io/en-US/), the backend is based on Python 3.10.15 implemented with Flask v2.0.3, and the whole is deployed on a Linux environment running Nginx. The entire platform operates within a Linux environment under Nginx, ensuring efficient and stable user interactions. This infrastructure supports seamless access to AIPAMPDS’s generation and screening functionalities.

## RESULTS

### Web server

The AIPAMPDS web server, accessible at https://amps.pianlab.cn/AIPAMPDS/, provides a freely available platform for generating and screening AMPs without requiring user registration or login, ensuring privacy and ease of access. Compatible with major web browsers and operating systems, it delivers consistent functionality across diverse devices. The AIPAMPDS home page offers a clear overview of its core functionalities: AMP generation via the ‘Generation’ tab, AMP screening via the ‘Screening’ tab, and a detailed tutorial via the ‘Help’ tab. Additionally, the ‘Download’ tab enables users to obtain the source code for local customization and execution (**Fig. 1D**). The following subsections detail the operation of the two primary features: AMP Generation and AMP Screening.

### AMP Generation

The AMP Generation tool focuses on creating novel and diverse AMP sequences through deep learning models. The generation tool contains three types of generation models that are based on different data fine-tuning: broad-spectrum AMP generation model fine-tuned with *E. coli* and *S. aureus* MIC dataset, *E. coli* generation model fine-tuned with *E. coli* MIC dataset, and *S. aureus* generation model fine-tuned with *S. aureus* MIC dataset. The users can determine the type of candidate AMPs to be generated by clicking the corresponding radio button to choose arbitrarily between the three models of broad-spectrum, *E. coli* and *S. aureus* (**Fig. 2A**). In addition, the user can set the generation parameters, including the number and maximum length of generated sequences. If the users do not enter any of the above, click ‘Submit’ directly, and the platform will generate 50 broad-spectrum AMPs with a maximum length of 50 by default for the user’s reference. Users can choose to fill in their personal email address and task name, and the system will send an email as a reminder after the task is completed. The web page output (**Fig. 2B**) shows the first 30 candidate AMP sequences generated by the model and the corresponding length, molecular weight, charge, instability index and other physicochemical properties. Users can also download the complete result file in CSV format for use.

**Figure 2.**
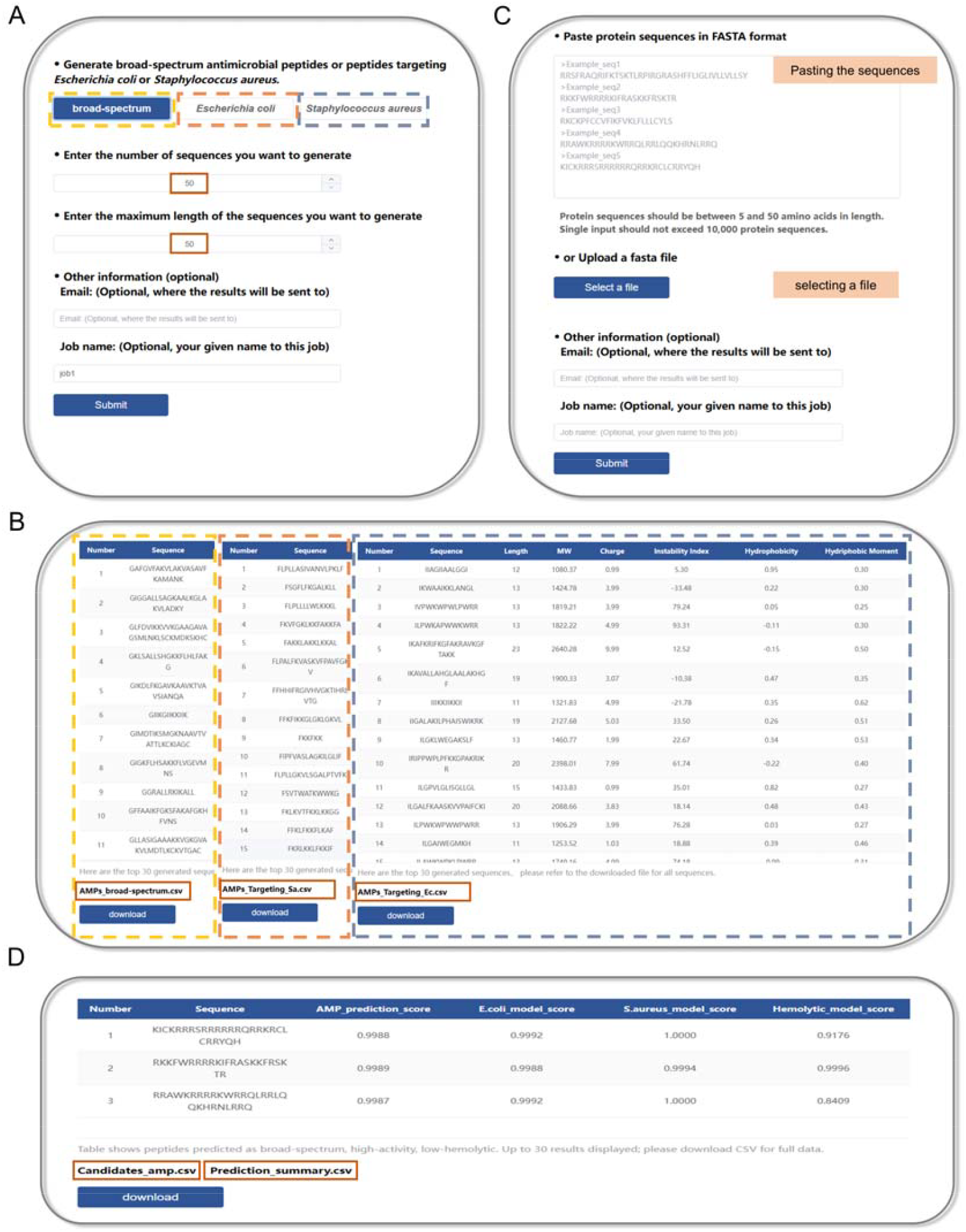
Introduction and usage of AIPAMPDS web server. **(A)** The AMP generation page of AIPAMPDS allows users to specify the target for AMP generation, set the number of generated sequences, and define the maximum sequence length. **(B)** The AMP generation results page displays the outcomes for three targets, with the results table containing the generated sequences and their corresponding physicochemical properties. Users have the option to download the complete results as a CSV document. **(C)** The AMP screening page of AIPAMPDS allows users to submit sequences through two methods, including pasting the sequences into the window or selecting a file from a local document. **(D)** The AMP screening results page presents the sequences of AMPs predicted by the model to have broad-spectrum high activity and low hemolytic activity, along with the model’s prediction scores. Users are able to download all results as a CSV file.

### AMPs Screening

The AMP Screening tool integrates four deep learning models for screening broad-spectrum, high-activity, low-hemolytic candidate AMPs from users-submitted candidate peptides. Users can also screen for antimicrobial peptides that meet specific requirements based on their needs. The tool accepts protein sequences in fasta file format as input data, which can be submitted by the user in one of two ways: either by pasting the peptide sequence in a text box or by uploading the fasta file to be analyzed (**Fig. 2C**). If the users do not submit the data in any way, also can click ‘Submit’ directly, and the platform will filter the sequences by default using the 5 sample sequences in the text box as inputs. Users can choose to fill in their personal email address and task name, and the system will send an email as a reminder when the task is completed. The screening page will return a table of the top 30 AMPs and the scores of each model (**Fig. 2D**). At the same time, the user can download two CSV files to view the specific results, including the model scores of all input sequences and the intersecting sequences and corresponding scores of the input sequences screened by the screening tool.

### Performance evaluation and comparison

To effectively identify AMPs meeting predefined criteria, we developed four deep learning models: AMP-Identifier, EcoliMIC-Classifier, StaphMIC-Classifier, and HemoRisk-Estimator. These models collectively assess antimicrobial activity, MIC against *E. coli* and *S. aureus*, and hemolytic potential, ensuring the selection of optimal AMP candidates. The performance of these models is evaluated and compared below.

We contrasted AMP-Identifier with state-of-the-art methods, including Ma *et al*. (30), amPEP (55), Macrel (56), AMPlify (57), and AMPscanner (29), using the test dataset distinct from the training dataset. Performance was assessed across multiple metrics: ACC, SEN, F1-score, and AUC for both ROC **(Fig. 3A)** and PR curves **(Fig. 3B)**, When benchmarked against the existing AMP classification models above, all metrics demonstrated enhanced performance of AMP-Identifier **(Table S1)**. Our model achieved a ROC of 99% (improvement of 4.2–7.7%) and an accuracy of 99.18% (improvement of 14.32–27.82%) **(Fig. 3, Table S1)**. These results highlight AMP-Identifier’s effectiveness and potential for accurate AMP classification. Compared with models trained solely with simple encoding, our model with a protein language model (ProtBERT(37)) embedding demonstrated precision improvements.

**Figure 3.**
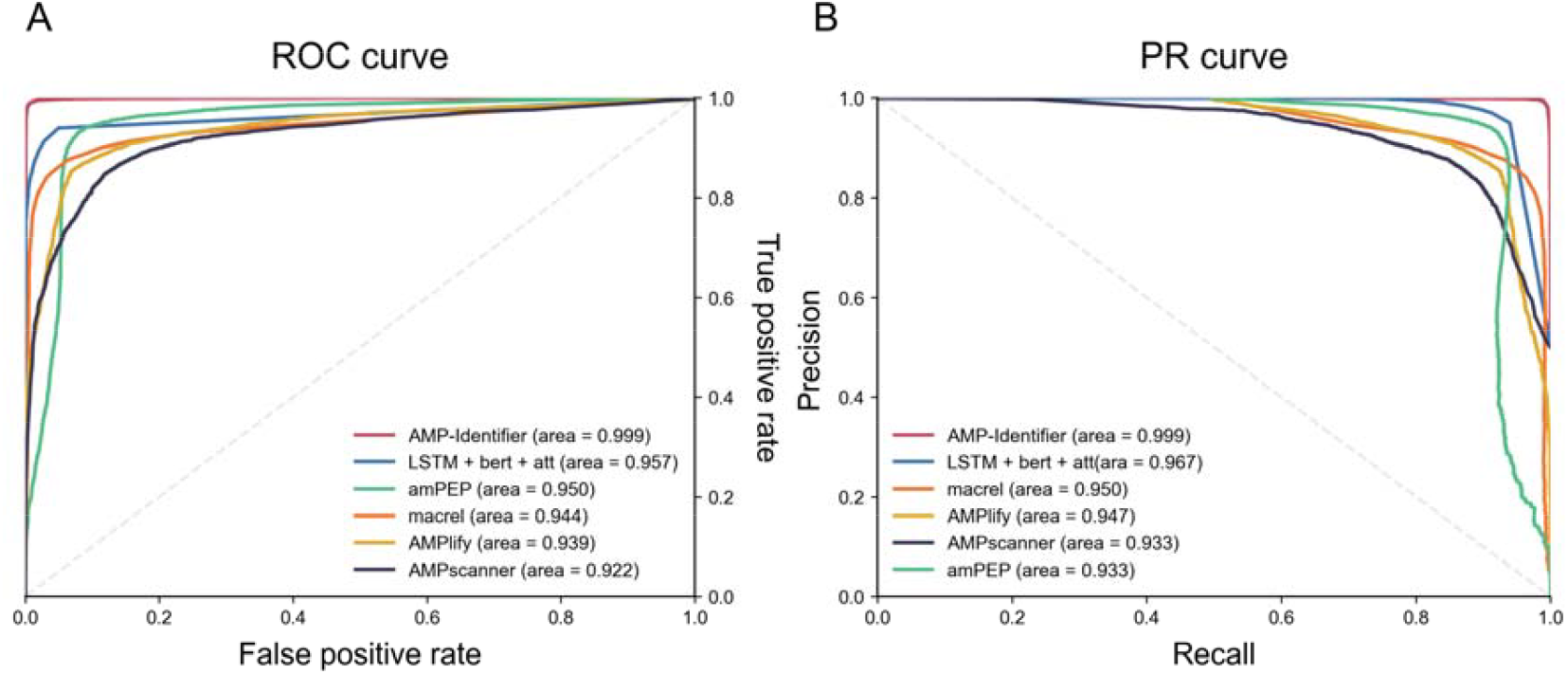
**(A)** Receiver operating characteristic curves of models in this study and previous methods. Precision–recall curves of models in this study and previous methods.

For the MIC prediction model, no comparison was made because other MIC models used regression models, which were different from our idea. We implemented 10-fold cross-validation of binary classification with EcoliMIC-Classifier and StaphMIC-Classifier to demonstrate the model’s potential to generalize to external data. **Table S2 and Table S3** show that the 10-fold cross-validation results indicate minimal impact of data partitioning on model performance. This evidence confirms the proposed MIC identification model’s efficiency in discovering high activity AMPs. The proposed model’s selection of an appropriate training model achieves a balance between performance and efficiency.

Because of the differences in the hemolysis data used by the individual models, we trained other models with the hemolysis data we collected, made predictions using an independent test set, and compared them afterwards. There were 2194 hemolytic and 1385 non-hemolytic peptide sequences used for training and validation. 244 hemolytic and 154 non-hemolytic peptide sequences as an independent test set. **Table 1** outlines the HemoRisk-Estimator we applied and other hemolysis models prediction accuracy. Comparison results show that the HemoRisk-Estimator model outperformed the other models in the hemolysis task, achieving an accuracy of 84.67%, while the embedding + Bi-LSTM and PeptideBERT approaches achieved 78.64% and 76.63% accuracies, respectively (**Table 1**). This showcases the robustness of our model in predicting hemolytic properties.

**Table 1.** Hemolysis classification accuracy comparison of previous methods and our HemoRisk-Estimator approach.

| Models | ACC | PRE | SEN | SPE | MCC | F1-score | AUC |
| --- | --- | --- | --- | --- | --- | --- | --- |
| HemoRisk-Estimator | **84.67** | **83.45** | **75.32** | 90.57 | **67.32** | **79.18** | **88.95** |
| PeptideBERT(58) | 76.63 | 74.02 | 61.039 | 86.48 | 49.65 | 66.9 | 83.44 |
| one-hots + RNN(59) | 76.38 | 68.29 | 72.73 | 78.69 | 50.88 | 70.44 | 83.51 |
| embedding + Bi-LSTM(60) | 78.64 | 81.65 | 57.79 | **91.80** | 54.17 | 67.68 | 84.27 |

### Validation

In order to validate the effectiveness of the generation and screening of the AI integrated computational framework employed by the AIPAMPDS platform, we carried out the validation in two ways: generate-screening process and natural candidate AMPs screening process.

For generate-screening process was as follows: we first generated 100,000 candidate AMPs using the AMP generation (*E. coli* option, targeting Gram-negative bacteria), and used these sequences as inputs to screen through the screening module to finally obtain 82 candidate AMPs. For natural candidate AMPs screening process was as follows: peptides collect from ruminant gastrointestinal tract metagenomes(61), and used these sequences as inputs to screen through the screening pipeline to finally obtain 9 candidate AMPs.

Considering that the structure of AMPs has a significant effect on their antimicrobial activity, we combined the secondary and tertiary structures of the candidate sequences for further screening. Firstly, the secondary structures of the sequences were calculated by DSSP software(62), and the protein sequences with 75% and above of α-helix were retained, and then the 3D structures of the candidate sequences were predicted by using AlphaFold2(63) (**Fig. S1**), and the sequences with loose structures were removed. After screening, five candidate AMPs (GAMPs) from the generated candidate AMPs were selected. The natural candidate AMPs were obtained by removing sequences greater than 30AAs in length, yielding five candidate AMPs (NAMPs) (**Fig. 1E**).

To assess the antimicrobial activity of the candidate AMPs (**Table S4**), ten candidates were synthesized and tested against representative Gram-positive (*S. aureus* ATCC 29213) and Gram-negative (*E. coli* ATCC 25922) bacteria. All natural candidate AMPs (NAMP_1, NAMP_2, NAMP_3, NAMP_4, NAMP_5) potent activity against both Gram-negative and Gram-positive bacteria (**Fig. 1E, Fig. 4A)**. For generated candidate AMPs, all of those are effective against Gram-negative bacteria, as the generation process specifically selects for options targeting Gram-negative bacteria. When extending to ESKAPE pathogens (*Enterococcus faecium, Staphylococcus aureus, Klebsiella pneumoniae, Acinetobacter baumannii, Pseudomonas aeruginosa*, and *Enterobacter* spp.), 90% of the candidate AMPs demonstrated antimicrobial effectiveness against the majority of these pathogens. In particular, GAMP_4 exhibits high antimicrobial activity against all ESKAPE pathogens (MICs: 8–64 μg/mL), demonstrating effective broad-spectrum antimicrobial activity (**Fig. 4A)**.

**Figure 4.**
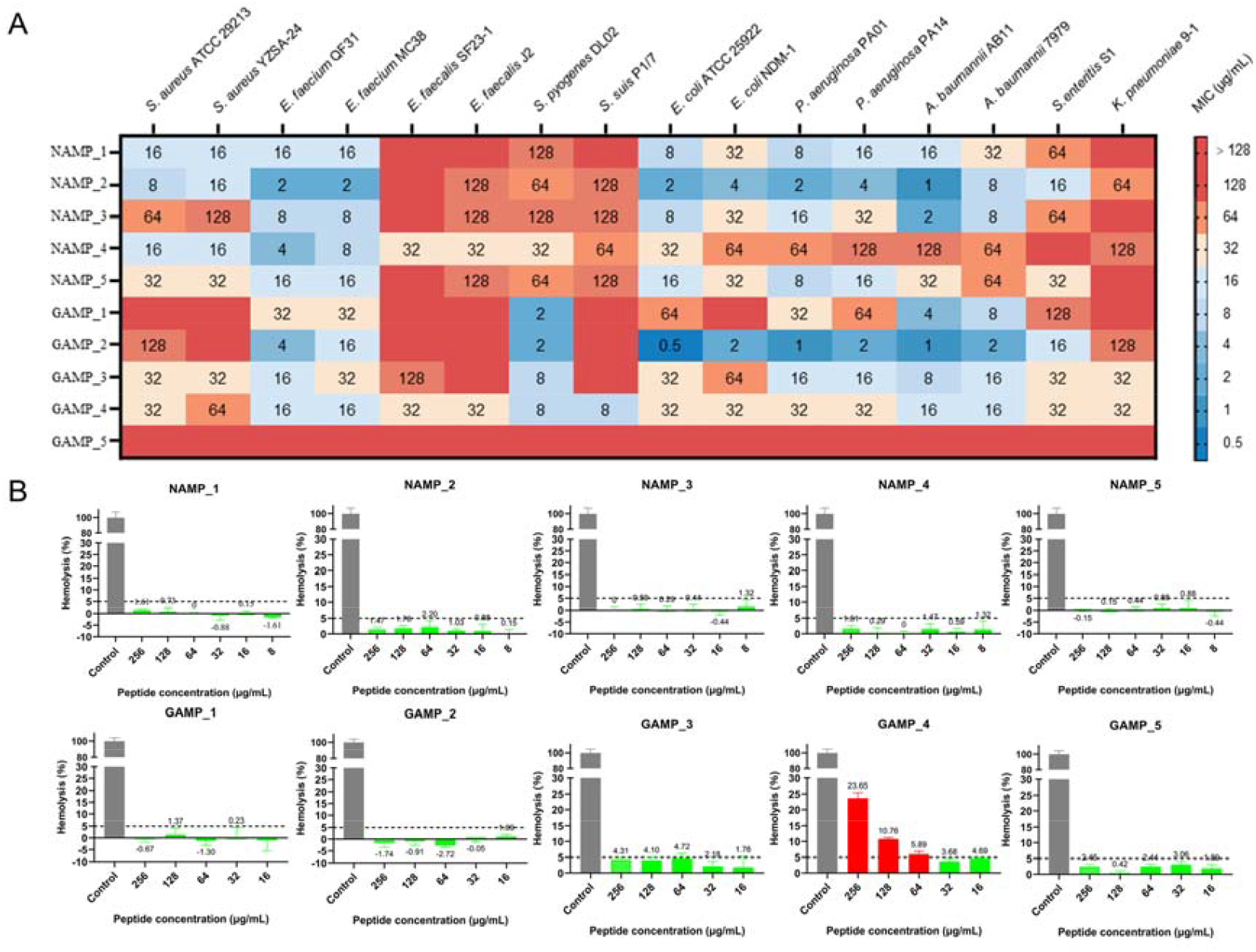
**(A)** MICs of ten candidate AMPs against diverse bacterial species. **(B)** Hemolytic toxicity of ten candidate AMPs on mice red blood cell.

In order to further evaluate the hemolytic toxicity of the nine candidate peptides, we further conducted hemolysis experiments on these peptides at different experimental concentrations (8-256 μg/ml) and counted the proportion of erythrocyte hemolysis caused by mice red blood cells (**Fig. 4B**). Hemolytic toxicity tests on mouse red blood cells showed that nine candidate AMPs displayed negligible hemolytic activity at the concentration of 128 µg/mL. In contrast, GAMP_4 caused 10.76% hemolysis the same concentration. These findings indicate that those candidate AMPs have minimal hemolytic toxicity, making it a safer candidate for therapeutic applications.

In conclusion, it can be seen that the AMPs generated by the AIPAMPDS platform and the peptides selected from natural AMPs by the screening pipeline showed excellent antimicrobial activity and good biocompatibility, which provides an effective tool and method for the development of AMPs.

## DISCUSSION

Although a significant number of methods have been developed for the discovery of novel AMPs(7,15,16,18), relatively few of these methods are capable of targeted generation and screening of AMPs. In this study, we introduce AIPAMPD, an AI-driven computational platform, designed for the conditional generation of AMPs and the screening of candidate AMPs with high activity and low hemolytic potential. In contrast to conventional methods, it facilitates controlled generation and screening, and is capable of managing multiple controls concurrently. It can be utilized for the generation and screening of broad-spectrum antimicrobial peptides, as well as for those specifically targeting Gram-negative or Gram-positive bacteria.

Furthermore, we have implemented numerous innovations in the generation model and screening models. The generative model is based on the GPT model architecture. The use of GPT for AMP generation mitigates issues associated with the instability of GAN training and the spatial distribution constraints of VAE models. The AMP-Identifier utilizes ProtBERT(37), a natural language processing-based model, for embedding, which enables the acquisition of more comprehensive sequence features. The model achieves an ROC of 99% on the test dataset, outperforming other models(29,30,55-57). The HemoRisk-Estimator employs CNN and LSTM network layers to extract both local and long-range contextual features from the input peptide sequences(64), demonstrating a superior capability in identifying hemolytic activity compared to existing models (58-60).

Using AIPAMPDS versatile toolbox, 90% of the candidate AMPs demonstrated efficacy against broad-spectrum strains and they were hypo hemolytic (**Fig. 4**). The experimental results validate the reliability of our AI platform. This has led to the idea of providing a user-friendly web interface to establish AIPAMPDS in everyday laboratory work. It can be used as the tool for generation and screening tasks related to AMP. The user interface is modern and user-friendly, with a choice of three generation modes in the generation interface and convenient download functionality in both the generation and filtering interfaces.

The two-sided approach of AIPAMPDS, the web interface and the API, also allows an easy migration from initial tests to user data projects in a kind of rapid prototyping. In the future, AIPAMPDS will continue supplementing more AMP generation function and AMP screening function. In addition, we will focus on expanding the generation of AMPs for different strains and improving the quality of screening, as well as providing users with more practical tools for AMP generation and screening in the future. With the development of peptide drugs peptide, tools like AIPAMPDS that offer ease of use and comprehensive functionality will be crucial in navigating the complexities of peptide data. The development of AIPAMPDS is a major advance in bioinformatics and sets the pace for future improvements in AMP drug development.

## DATA AVAILABILITY

AIPAMPD is available for free without registration: https://amps.pianlab.cn/AIPAMPDS/. The core scripts for the AIPAMPDS pipeline are available for free on github (https://github.com/LYRHeidi/AIPAMPDS).

## SUPPLEMENTARY DATA

Supplementary data are available at NAR Online.

## AUTHOR CONTRIBUTIONS

**Cong Pian**: Conceptualization, Writing-Review & Editing. **Jinhu Huang**: Conceptualization, Writing-Review & Editing. **Yong Yang**: Formal Analysis, Writing-Review & Editing. **Yanru Li**: Formal Analysis, Writing-Original Draft. **Liming, Gao**: Writing-Original Draft. **Wenqian Zhao**: Formal Analysis, Writing-Original Draft. **Ziyu Wang:** Formal Analysis. **Xianghan Xu**: Investigation. **Jinwen Ji:** Investigation. **Yuan Zhang**: Investigation. **Yufeng Liu**: Investigation. **Liangyun Zhang**: Writing-Review & Editing.

